# Multi-omics analysis identifies RFX7 targets involved in tumor suppression and neuronal processes

**DOI:** 10.1101/2022.12.05.519097

**Authors:** Katjana Schwab, Luis Coronel, Konstantin Riege, Erika K. Sacramento, Norman Rahnis, David Häckes, Emilio Cirri, Marco Groth, Steve Hoffmann, Martin Fischer

**Affiliations:** Computational Biology Group, Leibniz Institute on Aging – Fritz Lipmann Institute (FLI), Beutenbergstraße 11, 07745 Jena, Germany; Core Facility for Proteomics, Leibniz Institute on Aging – Fritz Lipmann Institute (FLI), Beutenbergstraße 11, 07745 Jena, Germany; Core Facility for DNA Sequencing, Leibniz Institute on Aging – Fritz Lipmann Institute (FLI), Beutenbergstraße 11, 07745 Jena, Germany

**Keywords:** RFX7, target genes, knock-out, p53, cancer research, neurological disorders

## Abstract

Recurrently mutated in lymphoid neoplasms, the transcription factor RFX7 is emerging as a tumor suppressor. Previous reports suggested that RFX7 may also have a role in neurological and metabolic disorders. We recently reported that RFX7 responds to p53 signaling and cellular stress. Furthermore, we found RFX7 target genes to be dysregulated in numerous cancer types also beyond the hematological system. However, our understanding of RFX7’s target gene network and its role in health and disease remains limited. Here, we generated RFX7 knock-out cells and employed a multi-omics approach integrating transcriptome, cistrome, and proteome data to obtain a more comprehensive picture of RFX7 targets. We identify novel target genes linked to RFX’s tumor suppressor function and underscoring its potential role in neurological disorders. Importantly, our data reveal RFX7 as a mechanistic link that enables the activation of these genes in response to p53 signaling.

## Introduction

The regulatory factor X 7 (RFX7) just recently emerged as a novel tumor suppressor in lymphoid cancers ^1^. It has been identified as a putative major cancer driver mutated in 13 to 15 % of Epstein-Barr Virus-negative Burkitt lymphoma ^2,3^, and CRISPR/Cas9-targeting of *Rfx7* in a mouse lymphoma model confirmed its function as a tumor suppressor ^4^. Moreover, *RFX7* alterations have been associated with chronic lymphocytic leukemia ^5–7^, diffuse large B cell lymphoma ^4,8^, and acute myeloid leukemia ^9^ in humans and with lymphoma ^4,10^ and leukemia ^11^ in mice. In addition to cancer, *RFX7* has been associated with metabolic disorders ^12^, neurological disorders ^13–15^, and organismal development and cellular differentiation ^16,17^.

The RFX transcription factor family is evolutionarily conserved and contains eight members in mammals that share a conserved DNA-binding domain recognizing X-box DNA motifs ^18–22^. Recently, we uncovered that RFX7 can be activated by the tumor suppressor p53 and cellular stress, such as DNA damage and ribosomal stress, in various cell types ^23^. Although p53 is one of the best studied proteins, the mechanisms underlying the p53-dependent regulation remain unknown for surprisingly many genes ^24–26^. The discovery of the p53-RFX7 signaling axis offered a mechanistic explanation for the p53-dependent regulation of multiple genes ^23^. Intriguingly, RFX7 target gene expression correlated with better patient prognosis across multiple cancer types, indicating a frequent dysregulation of RFX7 signaling in cancer even when RFX7 is not mutated ^23^. When we followed up on the RFX7 target DDIT4, we found that RFX7 inhibits AKT and mTORC1 activity both downstream and independent of p53 ^27^. Given that the inhibition of mTORC1 is an important means of p53 to suppress tumorigenesis ^28,29^, mTORC1 is likely to contribute also to the tumor suppressor function of RFX7 ^27^. Thus, the identification of RFX7 target genes is an important step to better understand the mechanisms contributing to RFX7’s role in the suppression of cancer and other diseases.

Here, we employed a multi-omics approach integrating transcriptome, cistrome, and proteome data from parental and RFX7 knock-out cells to identify additional RFX7 targets.

## Results

### Generation of RFX7 knock-out U2OS cells

We employed the osteosarcoma cell line U2OS which is frequently used to study the transcriptional program of p53 ^24^ because it contains functional p53 but amplified MDM2 ^30^.

First, we generated two RFX7 knock-out cell lines through CRISPR/Cas9 and single-cell cloning. Subsequently, we treated the parental and knock-out cell lines with the well-established MDM2 inhibitor Nutlin-3a to specifically activate p53 ^31^ and – if present – its downstream target RFX7 ^23^. Immunoblot analysis clearly confirmed the upregulation of p53 in all cell lines and the removal of RFX7 in the *RFX7*^-/-^ U2OS clones (**Figure 1a**). Specifically, RFX7 was present in DMSO control-treated parental U2OS cells and properly activated by Nutlin-3a treatment, as indicated by the lower migrating form of RFX7 ^23^.

**Figure 1.**
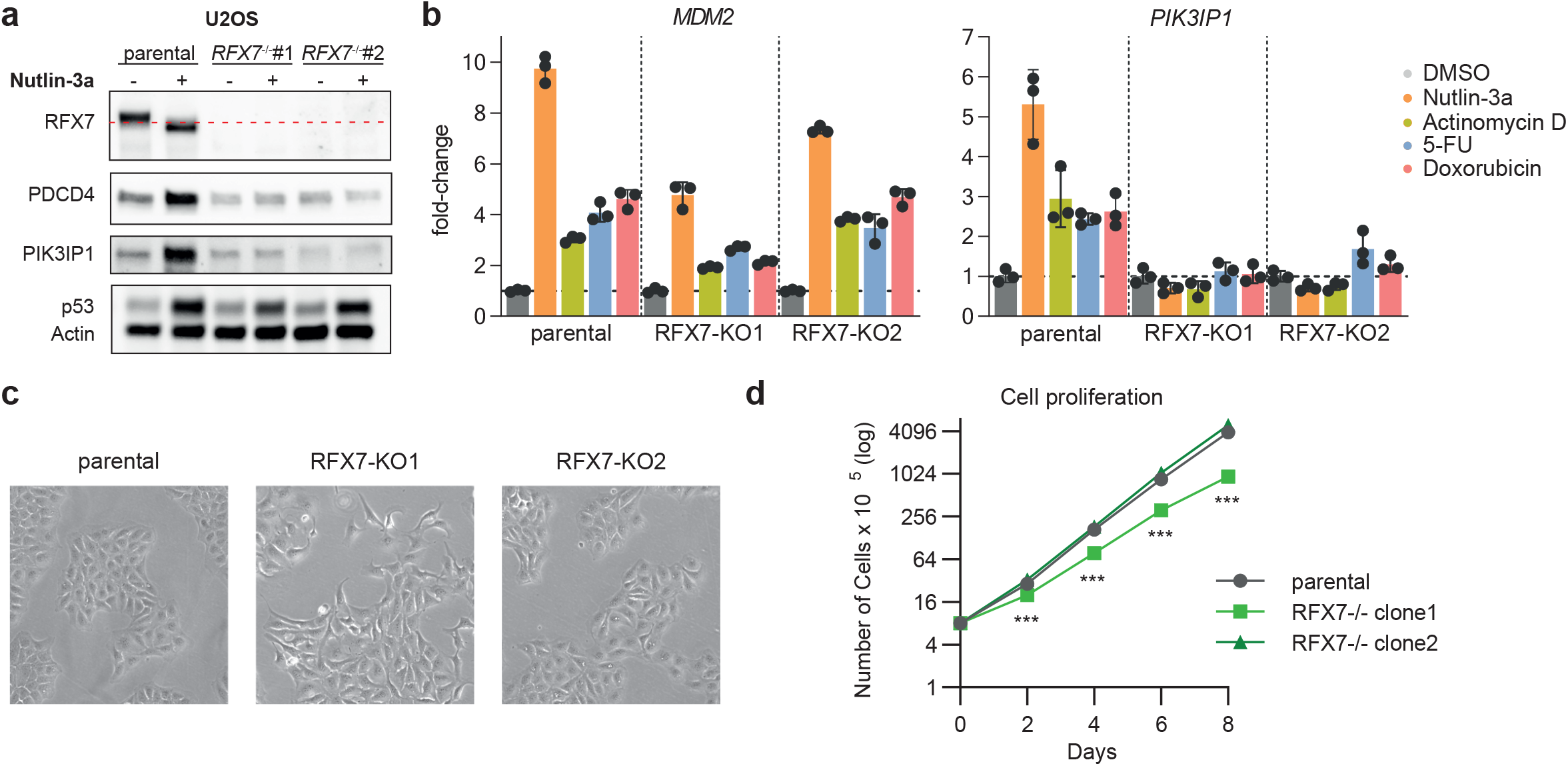
Generation and validation of U2OS RFX7 knock-out cell lines. **(a)** Western blot analysis of RFX7, PDCD4, PIK3IP1, p53, and actin (loading control) levels in parental and RFX7 knock-out (*RFX7*^*-/-*^) U2OS cells treated with Nutlin-3a or dimethyl sulfoxide (DMSO) solvent control. **(b)** RT-qPCR data of the RFX7 target *PIK3IP1* in in parental and RFX7 knock-out U2OS cells treated with Nutlin-3a, Actinomycin D, 5-FU, Doxorubicin, or DMSO solvent control. *MDM2* served as a positive control for p53 induction. Normalized to DMSO treatment and *ACTR10* negative control. Mean and standard deviation is displayed; n = 3 technical replicates. **(c)** Brightfield images of parental and RFX7 knock-out U2OS cells. **(d)** Cell number of parental and RFX7 knock-out U2OS cells following indicated days of cell culture.

Following up on our previous observation that *PDCD4* and *PIK3IP1* are directly induced by RFX7 ^23^, we found that PDCD4 and PIK3IP1 protein levels were up-regulated by Nutlin-3a in parental U2OS but not in the *RFX7*^-/-^ U2OS cell lines (**Figure 1a**). Using RT-qPCR we were able to confirm the RFX7-dependent gene regulation on the transcriptional level. To activate p53 signaling with other means than just Nutlin-3a, we also used the DNA-damaging agent Doxorubicin and the ribosomal stress inducers Actinomycin D and 5-FU. RT-qPCR analysis showed that the direct p53 target gene *MDM2*, serving as a positive control, was up-regulated in response to Nutlin-3a, Actinomycin D, 5-FU, and Doxorubicin in parental and RFX7 knock-out cells. In agreement with previous results ^23^, the RFX7 target gene *PIK3IP1* was up-regulated in response to Nutlin-3a, Actinomycin D, 5-FU, and Doxorubicin in parental U2OS cells. This up-regulation, however, was not observed in the two RFX7 knock-out clones, confirming the loss of functional RFX7 (**Figure 1b**). Notably, while the U2OS RFX7 knock-out clone #2 had an essentially undistinguishable growth rate and morphology compared with the parental U2OS cells, RFX7 knock-out clone #1 displayed somewhat altered morphology and slightly reduced proliferation (**Figure 1c** and **d**).

### Transcriptome analysis of RFX7 knock-out U2OS cells

To obtain a more comprehensive picture of the transcriptomic alterations in RFX7 knock-out cells, we performed Illumina sequencing of polyA-enriched RNA (RNA-seq) from parental U2OS and the two *RFX7*^-/-^ U2OS cell lines. Cells were treated with Nutlin-3a or DMSO control for 24 h. The transcriptome data reveals hundreds of differentially regulated genes in RFX7 knock-out cells compared with the parental U2OS cells (**Supplementary Table S1**). Comparing knock-outs and parental cells treated with Nutlin-3a, we observed the down-regulation of established direct RFX7 targets, such as *PIK3IP1, PDCD4, MXD4, RFX5, MAF, CCNG2, TSPYL1, PIK3R3* ^23^, and *DDIT4* ^27^ (**Figure 2a**). In agreement with a Nutlin-3a-mediated up-regulation of *PDCD4* and *PIK3IP1* through RFX7 ^23^, *PDCD4* and *PIK3IP1* induction by Nutlin-3a could not be observed in *RFX7*^-/-^ cells (**Figure 2b**). In fact, most of the 57 previously identified RFX7 target genes ^23^ were up-regulated in parental U2OS but not in the two knock-out cell lines in response to Nutlin-3a treatment. Moreover, RFX7 target gene expression was largely reduced in RFX7 knock-out cells compared with the parental cells also under control conditions (**Figure 2c**). These data further validate loss of functional RFX7 in the knock-out cell lines and corroborate the previously identified RFX7 target genes. Thus, our results validate the successful generation of two *RFX7*^-/-^ U2OS cell lines in which the p53-RFX7 signaling pathway is abrogated.

**Figure 2.**
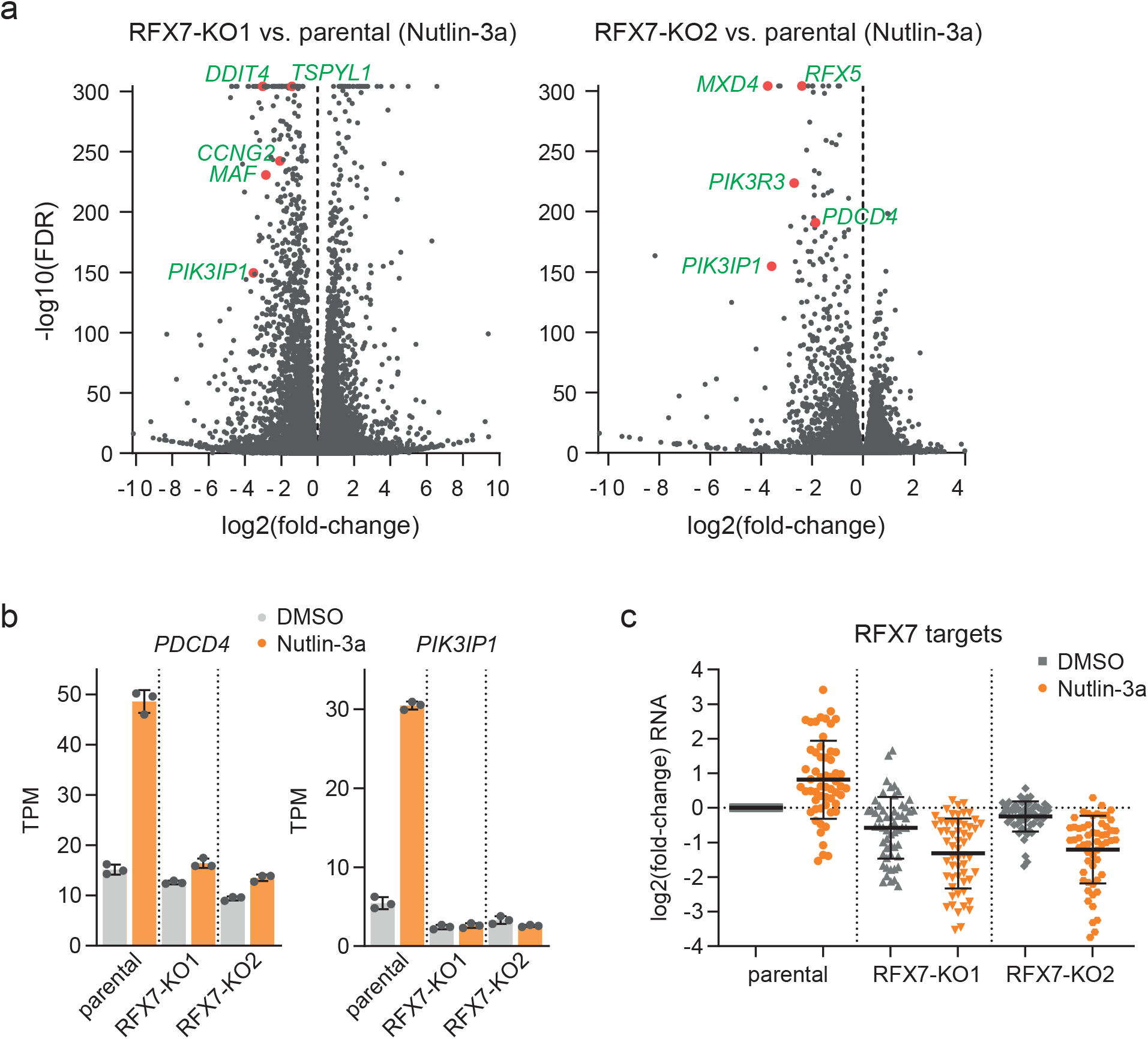
Transcriptome analysis of RFX7 knock-out U2OS cells. **(a)** Volcano plot of differential gene expression data from RFX7 knock-out clone 1 (left panel) and 2 (right panel) compared to parental U2OS cells under Nutlin-3a treatment condition. Data has been obtained using DESeq2 and is available in Supplementary Table S1. Selected known RFX7 targets are highlighted. **(b)** Transcripts per kilobase million (TPM) expression values of *PDCD4* and *PIK3IP1* obtained from RNA-seq analysis from parental and RFX7 knock-out U2OS cells treated with Nutlin-3a or DMSO solvent control. **(c)** Log2(fold-change) values of established RFX7 targets ^23^ from RNA-seq analyses of parental and RFX7 knock-out U2OS cells treated with Nutlin-3a or DMSO solvent control. Fold-changes normalized to DMSO-treated parental U2OS cells. Mean and standard deviation is indicated.

### Integrative analysis of transcriptome and cistrome data identifies novel RFX7 target genes

Since knock-outs often provide more robust signals compared to knock-down approaches, we asked whether transcriptome data derived from RFX7 knock-out cells may enable the identification of additional RFX7 target genes that were not identified in our previous RFX7 knock-down study ^23^. To this end, we integrated genome-wide DNA binding information derived from RFX7 ChIP-seq data from U2OS cells ^23^. We identified 33 genes that were not part of the previous list of 57 RFX7 target genes ^23^, but displayed an RFX7 binding site (‘peak’) near their transcription start site (TSS) and were significantly down-regulated in both RFX7 knock-out cell lines compared with parental U2OS under Nutlin-3a treatment condition (**Figure 3a** and **Table 1**). Many of the novel RFX7 targets became up-regulated by p53 (Nutlin-3a treatment), but this activation was largely absent in RFX7 knock-out cells. Thus, our present analysis strengthens the role of RFX7 as a critical mediator of p53-dependent gene regulation. For instance, *CLIC4* is long known to be p53-inducible, but the mechanism mediating *CLIC4* up-regulation in response to p53 signaling remained unresolved ^32^. Our data uncover *CLIC4* and many other genes to be regulated by the p53-RFX7 signaling axis (**Figure 3a**). Similar to the previous list of 57 RFX7 target genes ^23^ and in agreement with RFX7’s tumor suppressor function ^1^, we identified multiple tumor suppressors among the 33 novel targets, such as *MAFTRR* ^33^ (**Figure 2b**), *MIR22HG* ^34–36^, and *ARRDC3* ^37,38^ (**Figure 3c**). Moreover, supporting RFX7’s potential role in neuronal development ^16^ and neurological disorders ^13–15^, we identified regulators of neuronal processes among the novel RFX7 targets, such as *JUN* ^39,40^, *SYNPO* ^41–43^, *PPP3CA* ^44^, and *PPP2R5D* ^45–47^. The 33 novel RFX7 targets also include three anti-sense RNAs, namely *PDCD4-AS1, TOB1-AS1*, and *FAM111A-DT* (**Figure 3d**), expressed from gene loci overlapping previously identified RFX7 target genes. Intriguingly, three out of the 33 novel RFX7 target genes, namely *CCND2, JUN*, and *PNRC2*, are paralogs of the previously identified targets, i.e., *CCND1, JUNB*, and *PNRC1* ^23^. These data indicate that the respective ancestral genes may already have been under control of RFX7 (or its ancestor) and that the control mechanism has been conserved ever since.

**Figure 3.**
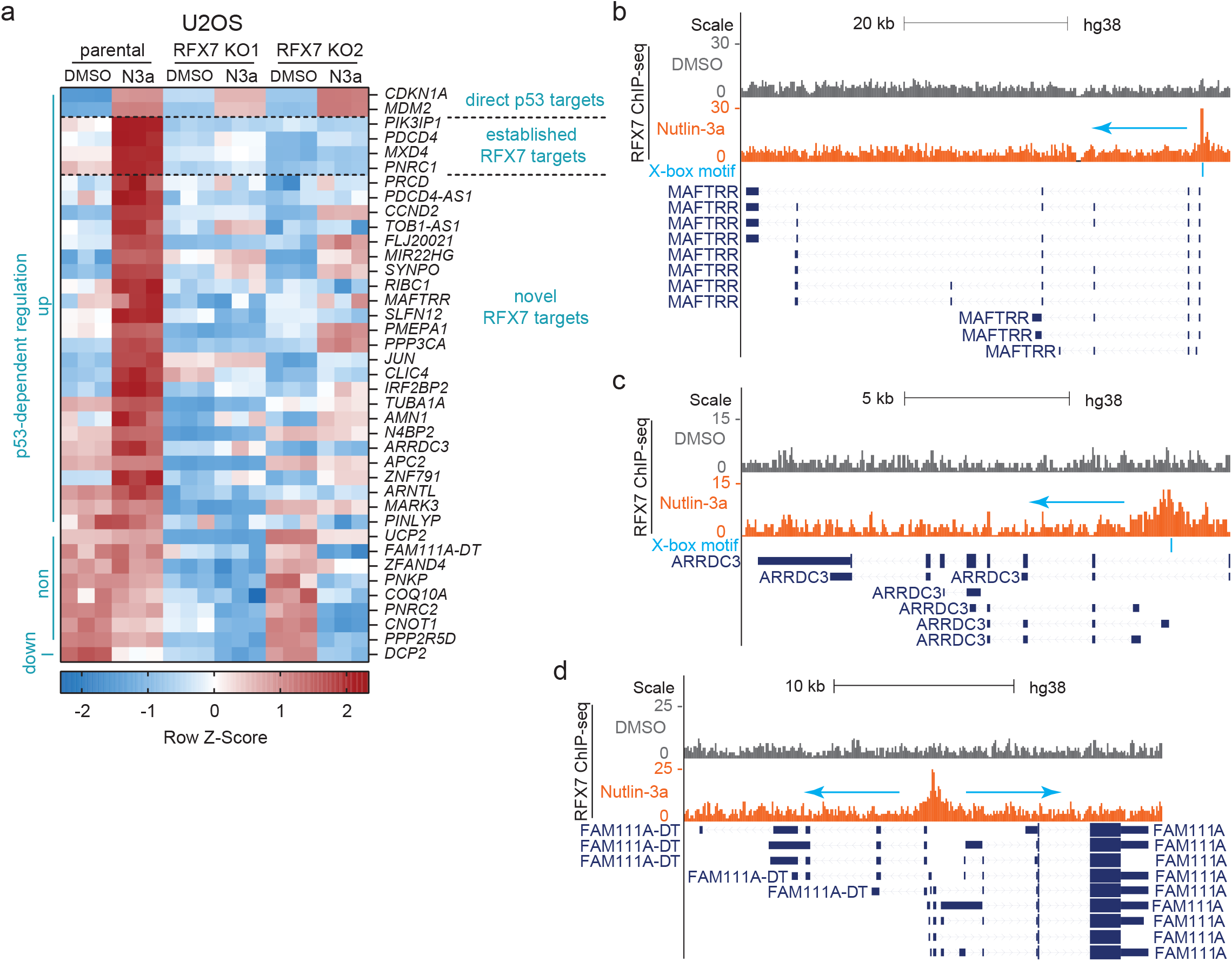
Integrative analysis data identifies novel RFX7 target genes. **(a)** Heatmap of RNA-seq data for established and novel direct RFX7 target genes that bind RFX7 within 5 kb from their TSS according to ChIP-seq data and are significantly (FDR < 0.01) down-regulated (log2FC ≤ –0.25) in both Nutlin-3a-treated RFX7 knock-out U2OS cell lines compared with parental U2OS cells. Significant (FDR < 0.01) p53-dependent regulation is indicated at the left. **(b-d)** Genome browser images displaying RFX7 ChIP-seq signals and predicted X-666 boxes at the **(b)** *MAFTRR*, **(c)** *ARRDC3*, and **(d)** *FAM111A-DT* gene loci.

Together, our transcriptome analysis of RFX7 knock-out U2OS cells combined with RFX7 DNA binding data revealed novel target genes contributing to RFX7’s roles as a tumor suppressor and potential neuronal regulator. Furthermore, the data provide a mechanism for the p53-dependent regulation of these genes and strengthens the role of RFX7 as a transductor of p53 signaling.

### Proteome analysis validates RFX7 targets

To complement the transcriptome data, we generated matched proteome data using mass-spectrometry analysis of whole cell lysates from ten biological replicates of Nutlin-3a and DMSO control-treated U2OS *RFX7*^*-/-*^ clone #2 and compared the data to proteome data from the parental U2OS cells. With proteomics, we quantified 5,392 unique proteins, including 19 out of the 57 known RFX7 targets and 11 out of the 33 novel RFX7 targets (**Supplementary Table S2**). The proteome data include the established RFX7 targets PDCD4 and ABAT known to be up-regulated by the p53-RFX7 signaling pathway ^23^ (**Figure 4a** and **b**). Another example is the cell cycle regulator CKS2, a target of RFX7 ^23^ as well as the *trans*-repressor complex DREAM that is activated through the p53 target p21 ^24,48^. In this way, p53-p21-DREAM mediated down-regulation of CKS2 is limited by p53-RFX7 signaling. Consequently, losing the RFX7 counterbalance leads to a more pronounced down-regulation or CKS2 (**Figure 4c**). Our proteomics analysis also confirmed the differential regulation of the novel targets JUN and SYNPO (**Figure 4d**). Overall, most of the known and novel RFX7 targets display reduced protein levels in RFX7 knock-out cells under Nutlin-3a treatment condition (**Figure 4e**). Together, the proteome analysis corroborates that proteins encoded by direct RFX7 targets largely follow the regulation of their encoding mRNA.

**Figure 4.**
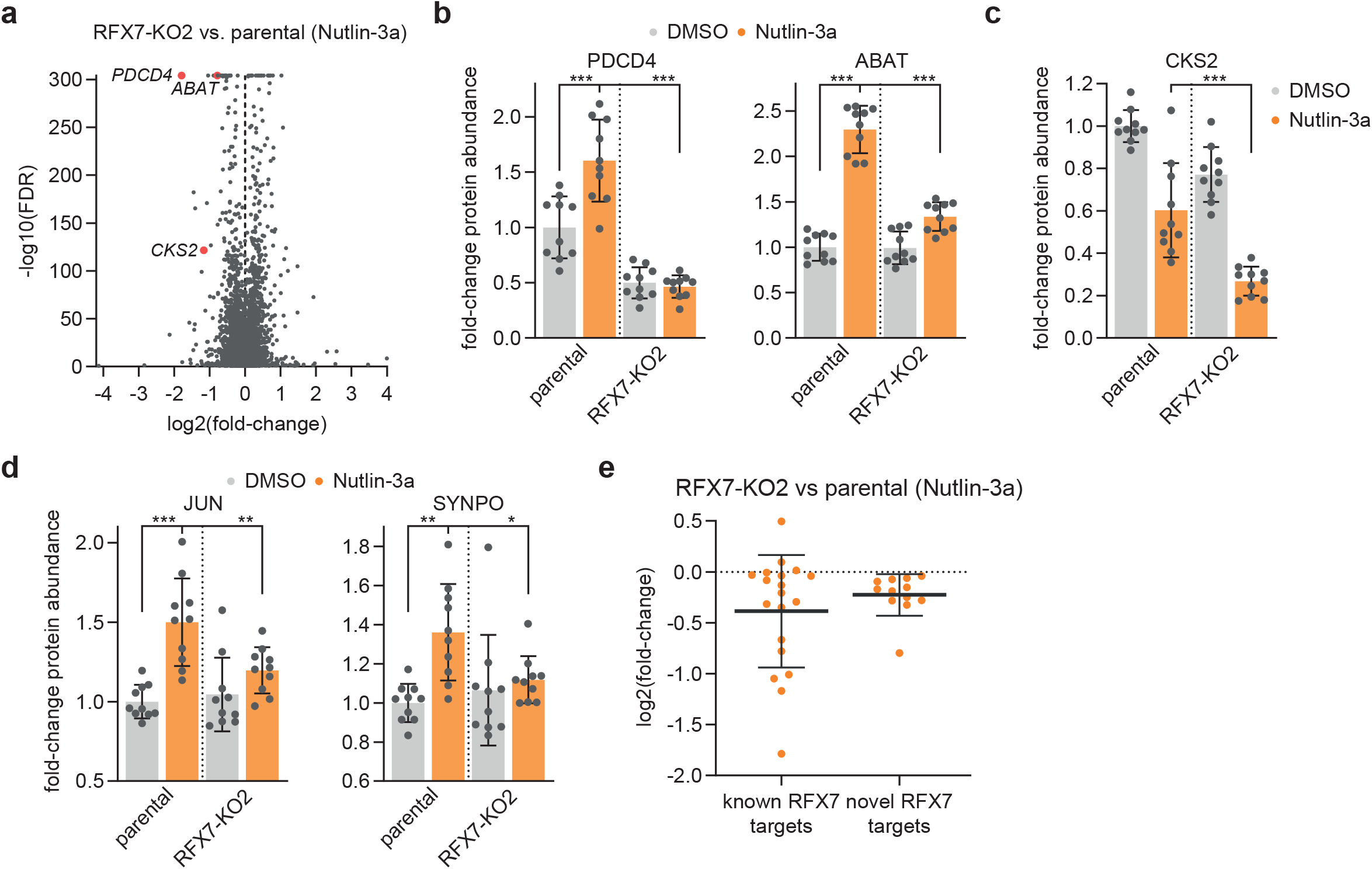
Proteome data validates RFX7 targets. **(a)** Volcano plot of differential protein expression data from RFX7 knock-out clone 2 compared to parental U2OS cells under Nutlin-3a treatment condition. Data has been obtained using Spectronaut and is available in Supplementary Table S2. Selected known RFX7 targets are highlighted. **(b-d)** Fold-change expression values of protein abundance. Normalized to DMSO-treated parental U2OS cells. Statistical significance obtained through a two-sided unpaired t-test, n = 10 biological replicates. **(b)** The established RFX7 targets PDCD4 (left) and ABAT (right). **(c)** The established RFX7 target, cell cycle gene, and DREAM target CKS2. **(d)** The novel RFX7 targets JUN (left) and SYNPO (right). **(e)** Log2(fold-change) values for 19 out of 57 established RFX7 targets ^23^ and 11 out of 33 novel RFX7 targets (Figure 3a) from mass spectrometry analyses of RFX7 knock-out U2OS cells compared with parental U2OS cells under Nutlin-3a treatment condition. Mean and standard deviation is displayed.

### Novel RFX7 targets are not specific to U2OS cells

To test whether the novel RFX7 targets are specific to U2OS or instead ubiquitously regulated by RFX7, we generated additional RFX7 knock-out clones from the non-cancerous retinal pigment epithelial (RPE-1) cell line via CRISPR/Cas9 and single-cell cloning. Again, we validated the removal of RFX7 by immunoblot analyses. Nutlin-3a increased p53 levels in all our RPE-1 cell lines and activated RFX7 in the parental RPE-1 cells (**Figure 5a**). We identified two *RFX7*^-/-^ RPE-1 clones, namely clones #3 and #27, in which RFX7 was no longer detectable. Similar to the U2OS cells (**Figure 1a**), PDCD4 and PIK3IP1 protein levels were up-regulated by Nutlin-3a in parental RPE-1 but not in the *RFX7*^-/-^ RPE-1 cell lines (**Figure 5a**). In addition to protein levels, we confirmed the loss of RFX7-dependent gene regulation using RT-qPCR. Similar to the U2OS cells, the RFX7 target genes *PIK3IP1, PDCD4, MXD4*, and *PNRC1* became up-regulated in response to Nutlin-3a in parental RPE-1 cells but not in the two RFX7 knock-out clones, confirming the loss of functional RFX7 (**Figure 5b**). We selected three novel RFX7 target genes, namely *MAFTRR, MIR22HG*, and *SYNPO*, for validation in RPE-1 cells. RT-qPCR data revealed that these novel RFX7 targets display a significantly reduced expression in Nutlin-3a-treated RPE-1 RFX7^-/-^ cells compared with parental cells (**Figure 5c**), suggesting that RFX7 regulates these genes across multiple cell types.

**Figure 5.**
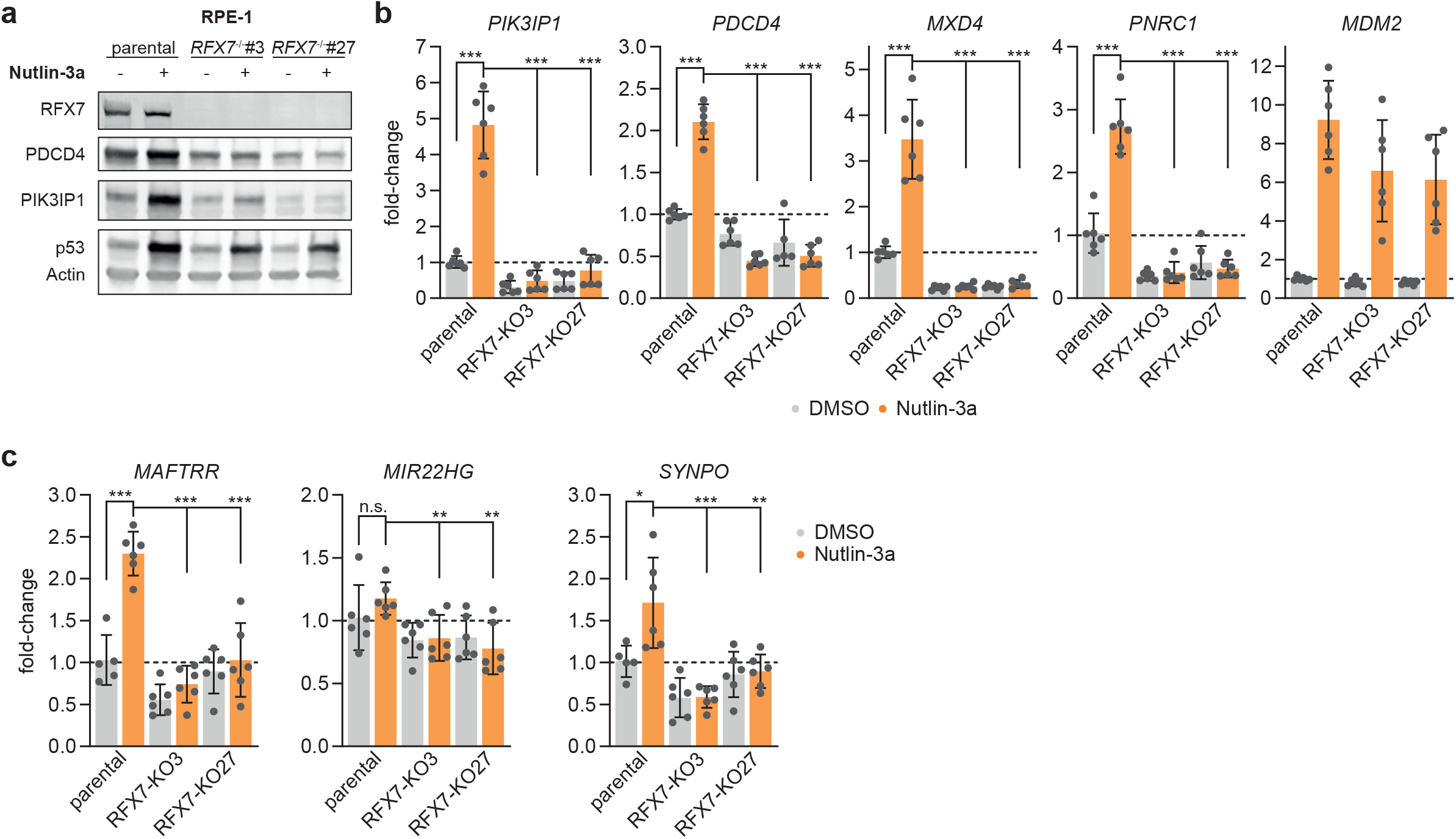
Generation of RPE-1 RFX7 knock-out cell lines and validation of novel RFX7 targets. **(a)** Western blot analysis of RFX7, PDCD4, PIK3IP1, p53, and actin (loading control) levels in parental and RFX7 knock-out (*RFX7*^*-/-*^) RPE-1 cells treated with Nutlin-3a or DMSO solvent control. **(b)** RT-qPCR data of the established RFX7 targets *PIK3IP1, PDCD4, MXD4*, and *PNRC1* in parental and RFX7 knock-out RPE-1 cells treated with Nutlin-3a or DMSO solvent control. *MDM2* served as a positive control for p53 induction. **(c)** RT-qPCR data of the novel RFX7 targets *MAFTRR, MIR22HG*, and *SYNPO* in parental and RFX7 knock-out RPE-1 cells treated with Nutlin-3a or DMSO solvent control. **(b and c)** Normalized to DMSO treatment and *ACTR10* negative control. Mean and standard deviation is displayed. Statistical significance obtained through a two-sided unpaired t-test, n = 6 (two biological replicates with three technical replicates each).

## Discussion

Recent research indicates that the understudied transcription factor RFX7 may have a role in tumor suppression ^1–4,23^, metabolic control ^12,17,27^, and neuronal fitness ^13–16^. However, it remains largely unclear how RFX7 elicits its functions. Thus, we have recently begun charting the gene regulatory network of the transcription factor RFX7 and identified 57 target genes ^23^. The study was critical to uncover the involvement of RFX7 in inhibiting the activity of the pro-survival kinases AKT and mTORC1 in part via its direct target DDIT4 ^27^. However, given the rather small number of target genes found using siRNA-mediated RFX7 knock-downs, we wondered whether additional target genes could be uncovered using a different strategy. Knock-out cell lines (**Figure 1a and 5a)** often provide more robust changes in signaling responses. Indeed, our multi-omics analysis of RFX7 knock-out U2OS cells identified 33 novel RFX7 target genes (**Figure 3a**), with validations on the proteome level (**Figure 4**) and in RFX7 knock-out RPE-1 cells (**Figure 5**). Many of the genes were up-regulated by p53, and RFX7 appears to be the key mediator of their p53-dependent up-regulation (**Figure 3a**).

The 33 novel RFX7 targets comprise multiple genes with known roles in tumor suppression or neuronal processes. For instance, the long non-coding RNA *MAFTRR* was found to promote transcriptional down-regulation of the neighboring *MAF* gene ^49^, which encodes for the transcription factor MAF that has important roles in lymphoid cell maturation ^50^ but which can function as an oncogene ^51,52^. RFX7 also has an important role in lymphoid cell maturation ^17^. Interestingly, we identified also *MAF* as an RFX7 target ^23^ and by activating both, *MAFTRR* and *MAF*, RFX7 may balance *MAF* expression to promote lymphoid cell maturation but avoid oncogenic transformation. Given that RFX7 has been associated with Alzheimer’s disease ^13^ and cognitive function ^14,15^, a particularly interesting candidate target is *SNYPO* encoding for the actin filament regulator synaptopodin, which is important for synaptic plasticity ^41,42^ and can facilitate Tau protein degradation ^43^. Intriguingly, *SYNPO* mRNA expression levels were found to be low in neurons from persons with Alzheimer’s disease ^53^, while a meta-analysis of cognitive trajectories in advanced age identified higher SYNPO protein levels to be associated with cognitive stability ^54^. Similar to synaptopodin, the phosphatase PPP3CA and the phosphatase PP2A regulatory subunit PPP2R5D are encoded by novel RFX7 target genes and are important for cognitive fitness ^44–47^. Together, the novel RFX7 targets yield additional evidence for RFX7’s role in tumor suppression and neurological disorders and provide promising starting points to further investigate its function. Interestingly, the combined list of previously established ^23^ and novel RFX7 target genes (**Figure 3a**) reveals pairs of paralogous genes, namely *CCND1*/*CCND2, JUN*/*JUNB, PNRC1*/*PNRC2, TOB1*/*TOB2*, and *TSPYL1*/*TSPYL2*. RFX7’s control over paralogous genes suggests that this regulation may have been conserved since the duplication events. Together with the evolutionary conservation of the whole RFX family and RFX7 itself ^21^, these findings underscore the importance of RFX7 and its target gene network from an evolutionary perspective.

In summary, our study expanded the list of RFX7 targets to facilitate a better understanding of RFX7 function and it provides a mechanistic understanding of how p53 can control those genes.

## Methods

### Cell culture, drug treatment, and transfection

U2OS cells (ATCC, Manassas, Virginia, USA) were grown in high glucose Dulbecco’s modified Eagle’s media (DMEM) with pyruvate (Thermo Fisher Scientific, Darmstadt, Germany). RPE-1 hTERT cells (ATCC) were cultured in DMEM:F12 media (Thermo Fisher Scientific). Culture media were supplemented with 10% fetal bovine serum (FBS; Thermo Fisher Scientific) and penicillin/streptomycin (Thermo Fisher Scientific). Cell lines were tested twice a year for *Mycoplasma* contamination using the LookOut Detection Kit (Sigma), and all tests were negative.

Cells were treated with DMSO (0.2 %; Carl Roth, Karlsruhe, Germany), Nutlin-3a (10 µM; Sigma Aldrich, Darmstadt, Germany), Actinomycin D (5 nM; Cayman Chemicals, Ann Arbor, Michigan, USA), 5-FU (25 μg/ml, Cayman Chemicals), or Doxorubicin (0.2 µg/ml; Cayman Chemicals) for 24 h.

### Generation of RFX7 knockout cells

Parental U2OS and RPE-1 cells were seeded in 6-well plates and the next day transfected with RFX7 CRISPR/Cas9 KO Plasmid (#sc-408041, Santa Cruz Biotechnology) using Lipofectamine 3000 (Thermo Fisher Scientific). 24 h after transfection, single cells expressing GFP (expressed from the CRISPR plasmid) were sorted into 96-well plates through flow cytometry using a BD FACSAria Fusion (BD Biosciences). Single-cell clones were cultured and tested for successful knockout of RFX7.

### RNA extraction and reverse transcription semi-quantitative real-time PCR (RT-qPCR)

Total cellular RNA was extracted using the RNeasy Plus Mini Kit (Qiagen, Hilden, Germany) or the innuPREP RNA Mini Kit (Analytik Jena, Jena, Germany) following the manufacturer protocol. One-step reverse transcription and real-time PCR was performed with a Quantstudio 5 using Power SYBR Green RNA-to-CT 1-Step Kit (Thermo Fisher Scientific) following the manufacturer protocol. We used *ACTR10* as a suitable control gene that is not regulated by p53 but expressed across 20 gene expression profiling datasets ^24^. The following RT-qPCR primers were used: *ACTR10* (forward: TCAGTTCCGGAAGGTGTCTT, reverse: GGACGCTCATTATTCCCATC), *MDM2* (forward: TCGGGTCACTAGTGTGAACG, reverse: TGAACACAGCTGGGAAAATG), *PIK3IP1* (forward: CCTGGTGCTACGTCAGTGG, reverse: TTCAGACGCTTCCTGGATTT), *PDCD4* (forward: GGTGGGCCAGTTTATTGCTA, reverse: GCACGGTAGCCTTATCCAGA), *MXD4* (forward: AAGCACAGACGAGCCAAACT, reverse: CCTGCTCCTCCAGTTTCTTG), *PNRC1* (forward: CGCATTTGAAGAAATCAGCA, reverse: CATCAGCTCCCTGTTTTGGT), *MAFTRR* (forward: CCTGGACAATGCTGGTTTTT, reverse: GCTGGTTTGAAGATGGAGGA), *MIR22HG* (forward: CTGAACTCCCTGGGAACAAG, reverse: TGAAGAACTACTGCGGCTCA), *SYNPO* (forward: GGGATCGAGGCTCAGGAC, reverse: GGCTCACCCAGCCGTCTA).

### Western blot analysis

Cells were lysed in Pierce IP lysis buffer (Thermo Fisher Scientific) containing protease and phosphatase inhibitor cocktail (Roche, Grenzach-Wyhlen, Germany or Thermo Fisher Scientific). Protein lysates were scraped against Eppendorf rack for 10 times and centrifuged with 15000 rpm for 15 min at 4°C. The protein concentration of supernatant lysates was determined using the Pierce 660 nm Protein Assay Kit (Thermo Fisher Scientific) and a NanoDrop ND1000 Spectrophotometer (Thermo Fisher Scientific). Proteins were separated in a Mini-Protean TGX 4-15% Gel (Bio-Rad) using Tris/Glycine/SDS running buffer (Bio-Rad). Proteins were transferred to a 0.2 µm or a low-fluorescence 0.45 µm polyvinylidene difluoride (PVDF) transfer membrane using a Trans-Blot Turbo (Bio-Rad). Following antibody incubation, membranes were developed using Clarity Max ECL (Bio-Rad) and a ChemiDoc MP imaging system (Bio-Rad) or, alternatively, ChemiDoc MP’s fluorescence detection was used.

Antibodies and their working concentrations: anti-mouse (1:5000; #7076, Cell Signaling Technology, or 1:20000, #STAR117D800GA, Bio-Rad), anti-rabbit (1:5000; #7074, Cell Signaling Technology or 1:10000, #12004162, Bio-Rad), actin (1:5000; #MA1-140, Thermo Fisher Scientific), RFX7 (1:1000; #A303-062A, Bethyl Laboratories), p53 (1:4000; kind gift from Bernhard Schlott ^55^), PDCD4 (1:1000; # 9535, Cell Signaling Technology), PIK3IP1 (1:500; #16826-1-AP, Proteintech).

### RNA-sequencing

Cellular RNA was extracted using the RNeasy Plus Mini Kit (Qiagen, Hilden, Germany) following the manufacturer’s protocol. U2OS cells were treated with Nutlin-3a to activate p53 signaling or with the DMSO solvent to serve as a negative control. Quality check and quantification of total RNA were performed using the Agilent Bioanalyzer 2100 in combination with the RNA 6000 Nano Kit (Agilent Technologies). Libraries were constructed from 500 ng total RNA using NEBNext Ultra II RNA - polyA+ (mRNA) Library Preparation Kit in combination with NEBNext Poly(A) mRNA Magnetic Isolation Module and NEBNext Multiplex Oligos for Illumina (96 Unique Dual Index Primer Pairs) following the manufacturer’s description (New England Biolabs).

Quantification and quality check of libraries were performed using a 4200 Tapestation instrument and D1000 ScreenTapes (Agilent Technologies). Libraries were pooled and sequenced on a NovaSeq 6000 (S1, 100 cycles). System run in 101 cycle/single-end/standard loading workflow mode. Sequence information was extracted in FastQ format using Illumina’s bcl2FastQ v2.20.0.422.

We used Trimmomatic ^56^ v0.39 (5nt sliding window approach, mean quality cutoff 22) for read quality trimming according to inspections made from FastQC (https://www.bioinformatics.babraham.ac.uk/projects/fastqc/) v0.11.9 reports. Illumina universal adapter as well as mono- and di-nucleotide content was clipped using Cutadapt v2.3 ^57^. Potential sequencing errors were detected and corrected using Rcorrector v1.0.3.1 ^58^. Ribosomal RNA (rRNA) transcripts were artificially depleted by read alignment against rRNA databases through SortMeRNA v2.1 ^59^. The preprocessed data was aligned to the reference genome hg38, retrieved along with its gene annotation from Ensembl v.102 ^60^, using the mapping software segemehl ^61,62^ v0.3.4 with adjusted accuracy (95%) and split-read option enabled. Mappings were filtered by Samtools v1.10 ^63^ for uniqueness and properly aligned mate pairs.

Following pre-processing of the data, read quantification was performed on exon level using featureCounts v1.6.4 ^64^, parametrized according to the strand specificity inferred through RSeQC v3.0.0 ^65^. Differential gene expression and its statistical significance was identified using DESeq2 v1.20.0 ^66^.

### Proteomics

Samples for proteomics analysis were prepared according to Buczak et al. ^67^. U2OS cells treated with Nutlin-3a or DMSO solvent control were sonicated using a Bioruptor (Diagenode, Beligum) for 10 cycles (60 seconds on and 30 seconds off with high intensity) at 20°C. The samples were then heated at 95°C for 10 minutes, before being subjected to another round of sonication. The lysates were clarified and debris precipitated by centrifugation at 14000 rpm for 10 minutes, then incubated with iodacetamide (room temperature, in the dark, 20 minutes, 15 mM). 10% of the sample was removed to check lysis on a Coomassie gel. Based on the gel, an estimated 25 µg of each sample was treated with 8 volumes ice cold acetone and left overnight at -20 °C to precipitate the proteins. The samples were centrifuged at 14000 rpm for 30 minutes, 4 °C. After removal of the supernatant, the precipitates were washed twice with 300 µL ice cold 80 % acetone solution. Each time after adding the acetone solution, the samples were vortexed and centrifuged again for 10 minutes at 4°C. The pellets were air-dried before being dissolved in digestion buffer at 1 µg/µL (1M guanidine HCl in 0.1M HEPES, pH 8). To facilitate the resuspension of the protein pellet, the samples were subjected to 5 cycles of sonication, as described above. Afterwards, LysC (Wako) was added at 1:100 (w/w) enzyme:protein ratio and digestion proceeded for 4 h at 37 °C under shaking (1000 rpm for 1 h, then 650 rpm). The samples were diluted 1:1 with milliQ water and were incubated with a 1:100 w/w amount of trypsin (Promega, sequencing grade) overnight at 37 °C, 650 rpm. The digests were then acidified with 10% trifluoroacetic acid and desalted with Waters Oasis® HLB µElution Plate 30µm under a slow vacuum following manufacturer instructions. Briefly, the columns were conditioned with 3×100 µL solvent B (80% acetonitrile; 0.05% formic acid) and equilibrated with 3x 100 µL solvent A (0.05% formic acid in milliQ water). The samples were loaded, washed 3 times with 100 µL solvent A, and then eluted into PCR tubes with 50 µL solvent B. The eluates were dried down with the speed vacuum centrifuge and dissolved in 5% acetonitrile, 95% milliQ water, with 0.1% formic acid at a concentration of 1µg/µL. 10 µL were transferred to an MS vial and 0.25 µL of HRM kit peptides (Biognosys, Zurich, Switzerland) was spiked into each sample prior to analysis by LC-MS/MS, which was performed according to Muntel at al. ^68^.

Peptides were separated on a nanoAcquity UPLC system (Waters) equipped with a trapping (nanoAcquity Symmetry C18, 5µm, 180 µm x 20 mm) and an analytical column (nanoAcquity BEH C18, 1.7µm, 75µm x 250mm). The outlet of the analytical column was coupled directly to Orbitrap Fusion Lumos (Thermo Fisher Scientific) using the Proxeon nanospray source. Solvent A was water, 0.1 % formic acid and solvent B was acetonitrile, 0.1 % formic acid. The samples (approx. 1 µg) were loaded with a constant flow of solvent A, at 5 µL/min onto the trapping column. Trapping time was 6 minutes. Peptides were eluted via a non-linear gradient from 1% to 40% B in 120 min. Total runtime was 145 min, including clean-up and column re-equilibration. The peptides were introduced into the mass spectrometer via a Pico-Tip Emitter 360 µm OD x 20 µm ID; 10 µm tip (New Objective) and a spray voltage of 2.2 kV was applied. The RF ion funnel was set to 30%.

For data independent acquisition (DIA) analysis, the conditions were as follows: Full scan MS spectra with mass range 350-1650 m/z were acquired in profile mode in the Orbitrap with resolution of 120,000 FHWM. The filling time was set at maximum of 20 ms with limitation of 5 × 10^5^ ions. DIA scans were acquired with 34 mass window segments of differing widths across the MS1 mass range. HCD fragmentation (normalized collision energy; 30%) was applied and MS/MS spectra were acquired with a resolution of 30,000 FHWM with a fixed first mass of 200 m/z after accumulation of 1 × 10^6^ ions or after filling time of 70 ms (whichever occurred first). Data were acquired in profile mode. For data acquisition and processing Tune version 2.1 and Xcalibur 4.0 were employed.

Acquired data were processed using Spectronaut Professional v13 (Biognosys AG). For library creation, the raw files were searched with Pulsar (Biognosys AG) against the human SwissProt database (Homo sapiens, entry only, release 2016_01) with a list of common contaminants appended, using default settings. For library generation, default BGS factory settings were used. DIA data were searched against this spectral library using BGS factory settings, except: Proteotypicity Filter = Only Protein Group Specific; Major Group Quantity = Median peptide quantity; Major Group Top N = OFF; Minor Group Quantity = Median precursor quantity; Minor Group Top N = OFF; Data Filtering = Qvalue; Normalization Strategy = Local normalization; Row Selection = Automatic.

### Statistics

RT-qPCR and expression data were analyzed using a two-sided unpaired t-test. Bar graphs display mean and standard deviation. *, **, ***, and n.s. indicate p-values <0.05, <0.01, <0.001, and >0.05, respectively. The number of replicates is indicated in each Figure legend. The experiments were not randomized and investigators were not blinded to allocation during experiments.

### Data availability

RNA-seq data are available through GEO ^69^ series accession number GSE173483. The mass spectrometry proteomics data are available through the ProteomeXchange Consortium via the Proteomics Identifications Database (PRIDE) partner repository ^70^ with the dataset identifier PXD024869. ChIP-seq track hubs are available through www.TargetGeneReg.org ^71^. Source data for Figures are available from the corresponding author upon request.

## Acknowledgements

This work was supported in part by the German Federal Ministry for Education and Research (BMBF) [031L016D to S.H] and the German Research Foundation (DFG) [FI 1993/6-1 to M.F. and HO 5281/7-1 to S.H.]. The FLI is a member of the Leibniz Association and is financially supported by the Federal Government of Germany and the State of Thuringia.

We thank Silke Förste for technical support and Bernhard Schlott for the kind gift of p53 antibody. We thank Ivonne Görlich from the FLI core facility DNA sequencing and Linda Rothenburger from the FLI core facility Flow Cytometry for diligent and skillful technical assistance.

## Author Contributions

M.F. conceived the study. M.F. and S.H. supervised the work. E.K.S. designed the mass spectrometry experiment. M.G. designed the RNA-seq experiment. M.F. designed the other experiments. N.R. performed the proteomics experiments. K.S., L.C., and D.H. performed the other experiments. E.C. analyzed the mass spectrometry data. K.R. analyzed the RNA-seq data. M.F. and K.S. analyzed the other data. M.F., S.H., and K.S. interpreted the data. M.F., with help from S.H. and K.S., wrote the manuscript. All authors read and approved the manuscript.

